# Genome-scale metabolic modeling and machine learning unravel metabolic reprogramming and mast cell role in lung cancer through a multi-level analysis

**DOI:** 10.1101/2025.03.06.641923

**Authors:** Masoud Tabibian, Tahereh Razmpour, Rajib Saha

## Abstract

Lung cancer remains a leading cause of cancer-related deaths worldwide, with immune interactions, particularly involving mast cells, playing a crucial role. Mast cells contribute to both pro- and anti-tumorigenic activities, influencing immune modulation, angiogenesis, and tissue remodeling. This study provides a comprehensive multi-level analysis of metabolic alterations in lung cancer through genome-scale metabolic modeling (GSM) and machine learning. Using 43 paired lung tissue samples, we developed metabolic models of lung cancer and mast cells, revealing a significant reduction in resting mast cells in cancerous tissues. Our Random Forest classifier accurately distinguished between healthy and cancerous states, identifying key metabolic signatures. We found that lung cancer cells selectively upregulate valine, isoleucine, histidine, and lysine metabolism in the aminoacyl-tRNA pathway to support their elevated energy demands. Mast cell metabolism exhibited enhanced histamine transport and increased glutamine consumption in the tumor microenvironment, suggesting a shift towards immunosuppressive activity. Additionally, our novel Metabolic Thermodynamic Sensitivity Analysis (MTSA) showed impaired biomass production in cancerous mast cells across physiological temperatures (36 to 40C), indicating metabolic vulnerabilities. By elucidating the metabolic adaptations of mast cells and lung cancer cells, our study highlights their interplay in tumor progression and identifies potential therapeutic targets and diagnostic markers for future investigation.

**Author Summary:** Our research examines the intricate relationship between lung cancer and mast cells, a type of immune cell, using advanced computational methods. We developed detailed metabolic models of lung tissue and mast cells through a multi-level approach to understand how their metabolism changes in cancer. Our findings reveal that lung cancer cells modify their metabolic pathways to meet increased energy demands, including enhanced utilization of four specific amino acids within the aminoacyl-tRNA pathway. In addition, mast cells in lung cancer environments show increased histamine release and glutamine consumption, suggesting they become more active in ways that might promote tumor growth. Additionally, we found that cancerous mast cells are less able to adapt to temperature changes compared to healthy ones, which could impact how they respond during fever or inflammation. These insights provide new perspectives on how lung cancer affects the immune system and could lead to novel approaches for diagnosis and treatment. By understanding these complex interactions, we aim to contribute to the development of more effective strategies for combating lung cancer.

## Introduction

Lung cancer remains one of the most prevalent and lethal cancers worldwide, accounting for a significant proportion of cancer-related deaths [1]. The identification of reliable biomarkers for lung cancer is crucial for improving early diagnosis and treatment strategies [2]. In particular, the complex interplay between cancer cells and the immune system has become a key focus in understanding lung cancer development and progression. The tumor microenvironment comprises various immune cells that significantly influence cancer progression, with mast cells playing a crucial role in this complex microenvironment[3].

Mast cells, traditionally known for their involvement in allergic reactions and inflammatory responses, have gained increasing attention in cancer research. These multifunctional immune cells are found in various tissues, including the lungs, and have been implicated in both pro-tumorigenic and anti-tumorigenic activities[3]. For instance, some studies have reported that a higher number of mast cells in non-small cell lung cancer (NSCLC) tissues are associated with poor patient prognosis[4]. Mast cells can contribute to anti-tumor immune responses through various mechanisms, including the promotion of dendritic cell maturation and migration, enhancement of CD8+ T cell activation and proliferation, and modulation of regulatory T cells (Tregs)[5]. However, mast cells can also exhibit pro-tumorigenic effects by promoting angiogenesis, tissue remodeling, and immunosuppression in certain contexts[6].

Computational methods can complement experimental efforts[7], which are often costly and constrained by limited data. Novel approaches that uncover metabolic signatures associated with lung cancer and mast cell activity could revolutionize diagnostic and prognostic capabilities in clinical settings. Genome-scale metabolic modeling (GSM) has emerged as a powerful tool in cancer research, enabling the comprehensive analysis of cellular metabolism at a systems level. By integrating genomic, transcriptomic, and metabolomic data, GSMs provide a comprehensive view of metabolic processes and their alterations in cancer cells[8]. These models have been successfully applied to various cancer types, such as pancreas [9], breast[10], colorectal[11], and brain[12]cancers, offering insights into metabolic vulnerabilities and potential therapeutic targets.

The field of cancer screening has been further enhanced by the integration of machine learning approaches. These computational techniques can identify complex patterns and relationships within large-scale datasets, facilitating the discovery of novel biomarkers and metabolic signatures associated with cancer progression[13]. Thus, the combination of GSM and machine learning holds great promise for advancing our understanding of cancer metabolism and improving diagnostic and therapeutic strategies.

This study aims to address a significant gap in the field by developing the first genome-scale metabolic models of lung cancer cells and mast cells in both healthy and cancerous states. By focusing on the metabolic role of mast cells in lung cancer, we seek to uncover novel insights into the complex interactions between these immune cells and the tumor microenvironment. Our objectives include the construction and analysis of GSMs for lung tissue and lung-associated mast cells, as well as the identification of specific metabolic biomarkers that could serve as potential diagnostic or therapeutic targets. Through this comprehensive approach, we aim to elucidate the metabolic reprogramming that occurs in mast cells within the context of lung cancer, potentially uncovering new avenues for targeted therapies and improved diagnostic strategies. By leveraging the power of GSM and machine learning techniques, this study represents a significant step forward in our understanding of the metabolic landscape of lung cancer and the role of mast cells in its development and progression.

## Results

### Cell Type Deconvolution Analysis

Understanding the complex cellular composition of the tumor microenvironment is crucial for unraveling the mechanisms of cancer progression. Traditional bulk RNA sequencing data, while informative, presents challenges in distinguishing signals from different cell populations within heterogeneous tissue samples. To address this limitation, we employed CIBERSORTx[14], a computational deconvolution method that enables accurate estimation of cell type proportions from bulk tissue transcriptomics data. This tool is particularly valuable for our study as it allows us to quantify the relative abundance of different immune cell populations, including mast cells, in both healthy and cancerous lung tissues.

CIBERSORTx analysis revealed significant differences in immune cell populations between healthy and cancerous lung tissues, with a particularly notable finding regarding resting mast cells (Fig 1). The analysis demonstrated a highly significant reduction in resting mast cell infiltration in cancerous lung tissues compared to healthy lung tissue (mean fraction ≈ 0.15 in healthy vs. ≈ 0.08 in cancer). The strong statistical significance of this finding is supported by both a very low p-value (1.315e-4) indicating the extreme unlikelihood that this difference occurred by chance, and a robust t-statistic (4.009) demonstrating that the observed difference is several standard deviations from zero. This substantial decrease suggests a potential shift in mast cell states during cancer development, raising important questions about their role in lung cancer pathogenesis. While our analysis also revealed significant alterations in other immune cell populations, including M2 macrophages (p = 4.598e-12, t = -8.060), dendritic cells (resting: p = 8.549e-13, t = 8.424; activated: p = 0.0448, t = -2.037), memory B cells (p = 1.926e-6, t = 5.118), and regulatory T cells (Tregs) (p = 7.526e-4, t = -3.498), this study focuses specifically on understanding the implications of reduced resting mast cells in the tumor microenvironment due to their multifaceted role in lung cancer. The significant decrease in resting mast cells, coupled with the complex remodeling of the immune landscape, provides a strong rationale for investigating their potential role in lung cancer progression or suppression.

**Fig 1.**
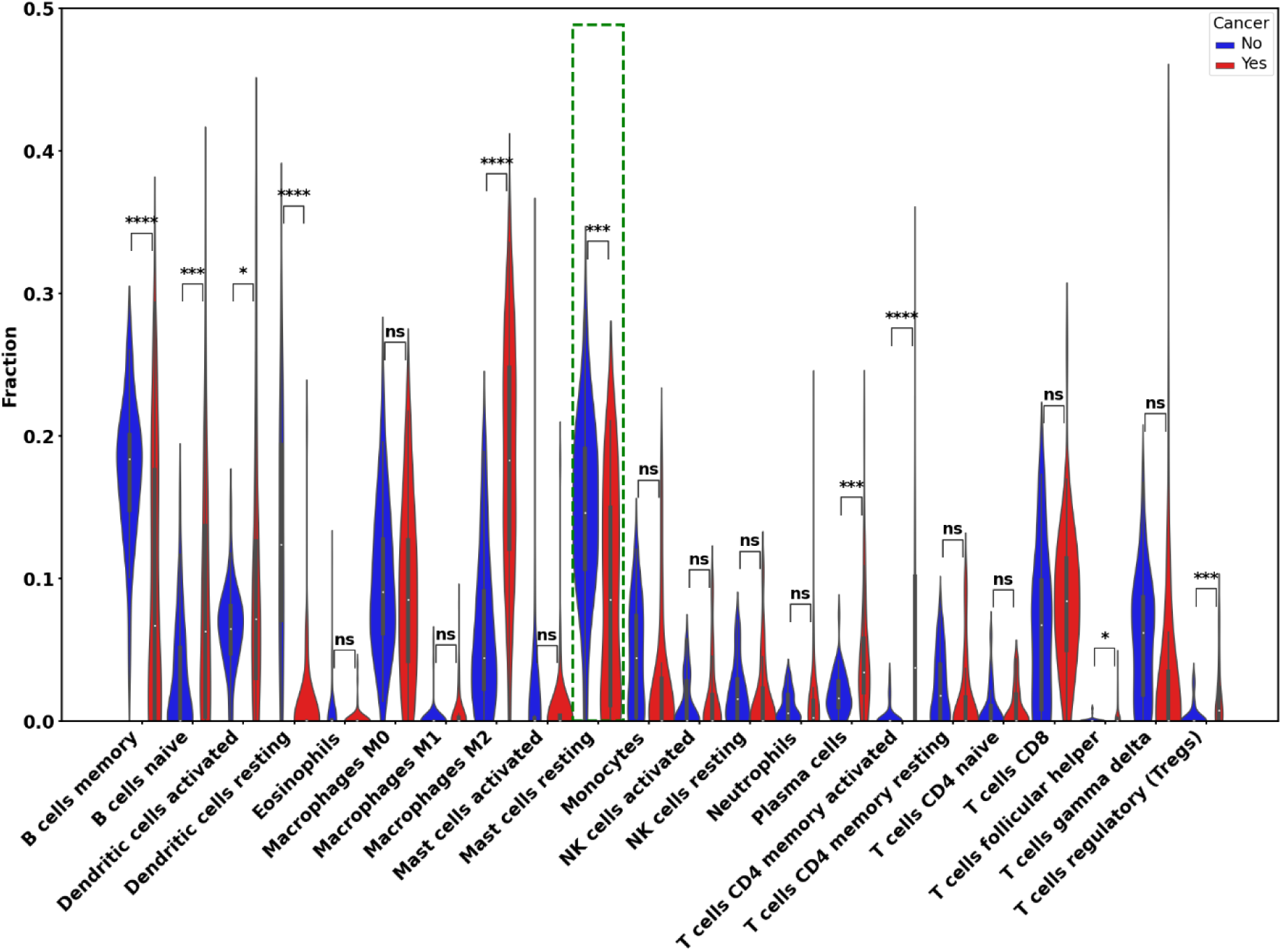
Immune cell composition analysis comparing healthy and cancerous lung tissue samples using the CIBERSORTx deconvolution method. The plots display the fractional abundance of 22 distinct immune cell populations, with blue representing healthy samples and red representing cancerous samples. Statistical significance between groups is indicated by asterisks (p≤0.0001**, *p≤0.001, p≤0.01, p≤0.05, ns: not significant). Notably, resting mast cells (highlighted by a green dashed box) show significant differences (p < 0.001) between healthy and cancerous conditions, suggesting their potential role in the lung cancer microenvironment. The y-axis represents the relative fraction of each cell type, and the width of each violin plot indicates the density distribution of the data points.

Based on these significant differences in immune cell populations, particularly the reduction in resting mast cells, we proceeded to develop genome-scale metabolic models to further investigate the metabolic changes associated with lung cancer and the role of mast cells in this disease.

### Genome-Scale Metabolic Modeling

We successfully developed genome-scale metabolic models for both bulk lung tissue and mast cell-specific samples using the Human1[15] model and implementing the iMAT[16] method. To validate our genome-scale metabolic models, we conducted in silico nutrient deprivation experiments by systematically analyzing the impact of essential nutrient sources (N, P, C, and S) in the models’ environmental conditions.

Our simulations demonstrated that carbon source elimination resulted in complete cessation of biomass production, as cells cannot synthesize carbon from scratch. Similarly, when all key nutrient sources were eliminated simultaneously, the models predicted complete growth arrest, aligning with the fundamental biological principle that cells cannot sustain growth without essential nutrients[17]. These outcomes support the models’ validity and adherence to basic metabolic constraints, confirming their biological accuracy and predictive capabilities. The genome-scale metabolic models generated for bulk lung tissue and mast cells exhibited varying numbers of reactions and metabolites across the 86 samples. Supplementary Figures (S1 Fig and S2 Fig) illustrate the distribution of these reactions and metabolites across all models. With these validated metabolic models, we next applied machine learning techniques to identify key metabolic differences between healthy and cancerous states.

### Machine Learning Analysis

#### Random Forest Classification

Leveraging the genome-scale metabolic models developed, we employed a Random Forest classifier to distinguish between healthy and cancerous states using reaction flux features derived from flux balance analaysis (FBA). The classifier achieved robust performance metrics for bulk lung tissue samples, with a mean accuracy of 0.95. The model exhibited balanced precision and recall scores of 0.95 and 0.94, respectively, resulting in an F1-score of 0.94. The confusion matrix reveals effective classification with minimal misclassification between healthy and cancer states (S1 Table in supporting information). For the mast cell models, the classifier achieved perfect classification metrics across all evaluation parameters, with accuracy, precision, recall, and F1-score all reaching 1.0. This exceptional performance can be attributed to the minimal variation in reaction fluxes between samples, as determined through FBA using gene expression data obtained from CIBERSORTx deconvolution. The confusion matrix demonstrates flawless separation between healthy and cancerous states, with zero misclassifications in both categories.

### Multi-level analysis reveals complex metabolic reprogramming in lung cancer

Our genome-scale metabolic modeling and Random Forest classification framework provide a powerful, multi-faceted lens through which to examine metabolic reprogramming in lung cancer. This integrated approach achieved robust differentiation between healthy and cancerous states with a mean accuracy of 0.95, while simultaneously revealing the underlying metabolic features driving this distinction. By combining reaction scores obtained through feature selection and deriving gene importances using Gene-Protein-Reaction (GPR) relationships; while also calculating pathway importance based on the reactions present in each specific pathway, we gained comprehensive insights spanning from pathway-level activities to individual gene contributions.

The Fig 2a highlights the top 40 reactions ranked by their importance in distinguishing between healthy and cancerous states, as determined through feature selection analysis in the first level of our multi-level metabolic alteration analysis in lung cancer. Among these, a reaction involved in the production of *benzoate* (MAR08643) is identified as the highest-ranked reaction. The large difference in flux values between healthy and cancer samples (S3 Fig in supporting information files) is a key reason for this reaction’s top ranking. In healthy samples, this reaction exhibits consistently high flux values, whereas in cancer samples, the flux is predominantly near zero, indicating that this reaction is mostly inactive in the cancerous state.

**Fig 2.**
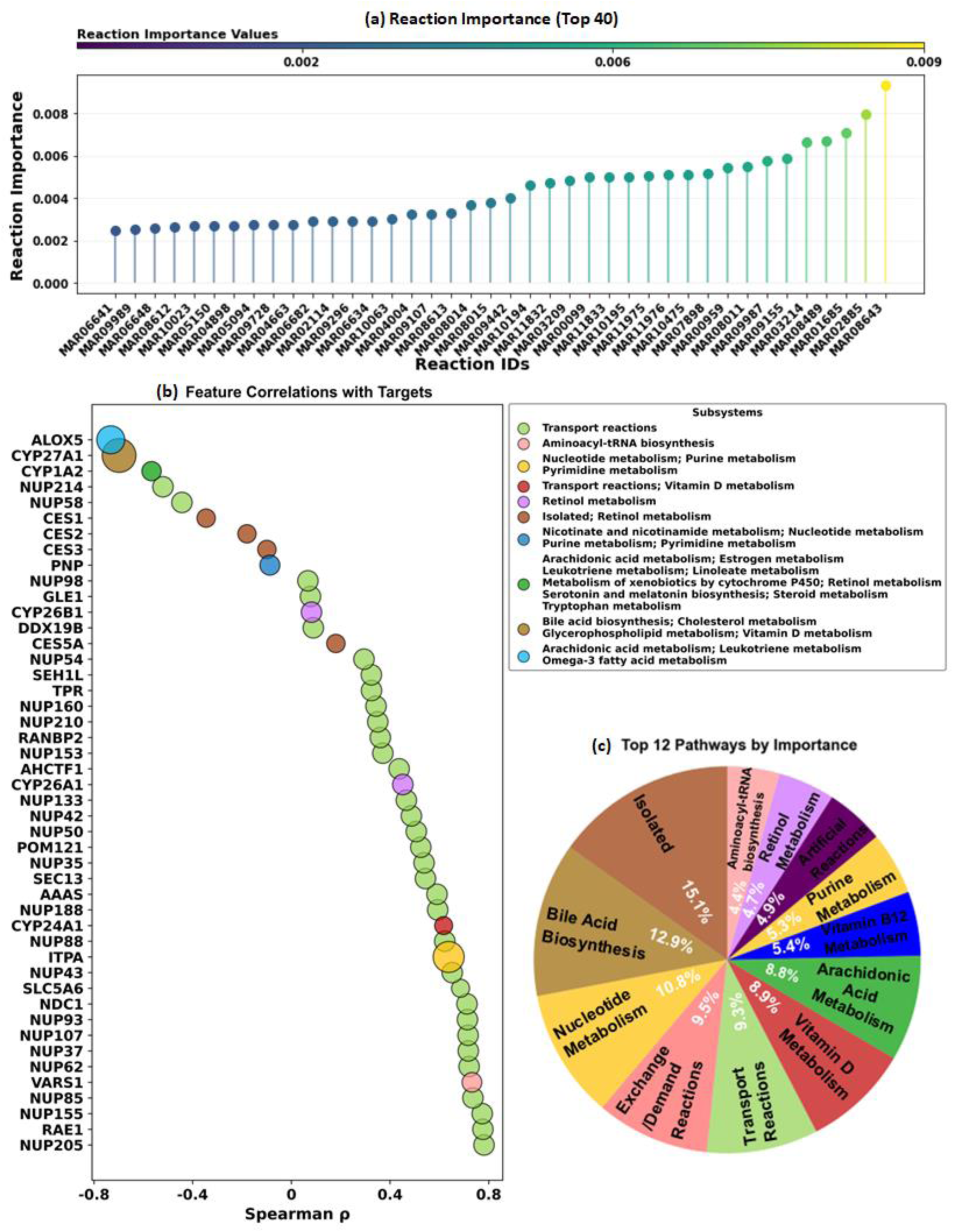
Multi-level Analysis of Metabolic Alterations in Lung Cancer: Gene, Reaction, and Pathway Level Results. (a). This plot displays the top 40 most important reactions in lung cancer metabolism. The reactions are ranked by their importance values (y-axis), with reaction IDs shown on the x-axis. A color gradient visually enhances the importance hierarchy. (b). The bubble plot illustrates the relationship between gene importance and their correlation with cancer state. The x-axis represents Spearman correlation coefficients (ρ), where positive values indicate upregulation and negative values indicate downregulation in cancer. Bubble sizes correspond to the relative importance of each gene, with larger bubbles indicating higher importance. Colors denote different metabolic subsystems, revealing pathway-level patterns. Notable features include CYP27A1, showing high importance (large bubble size) and strong negative correlation in bile acid biosynthesis pathway (purple), and several transport-related genes (pink) with positive correlations. The legend provides a comprehensive mapping of metabolic subsystems to their respective colors, highlighting the diverse metabolic processes involved in lung cancer metabolism.(c). Relative importance distribution of the top 12 metabolic pathways identified in lung cancer. The pie chart shows the percentage contribution of each pathway, with Isolated subsystem (15.1%), Bile acid biosynthesis (12.9%), and Nucleotide metabolism (10.8%) emerging as the most significant pathways.

The near-complete suppression of this reaction in cancer samples suggests that it is downregulated as part of metabolic reprogramming in lung cancer. This downregulation may reflect a strategy by lung cancer cells to suppress reactions involved in benzoate production. Benzoate has been implicated in modulating inflammatory responses through NF-κB activation, a pathway that plays a dual role in cancer biology. While NF-κB activation can promote tumor progression by supporting cell survival, angiogenesis, and metastasis, excessive inflammation can attract immune cells capable of attacking tumor cells [18]. By reducing flux through this reaction and limiting benzoate production, lung cancer cells may avoid triggering pro-inflammatory responses that could disrupt their immunosuppressive microenvironment or harm their survival. Other notable reactions include those involved in bile acid biosynthesis, such as MAR03214 and MAR01685, which will be discussed in detail in subsequent sections.

In the next level of our analysis, we examined gene-level correlations with lung cancer metabolic reprogramming. Our correlation analysis, visualized in Fig 2b, revealed distinct patterns of both positive and negative associations, with Spearman correlations ranging from -0.8 to 0.8. The relative importance of each gene, represented by bubble size, highlighted key players in metabolic reprogramming. Most notably, sterol 27-hydroxylase ,CYP27A1, emerged as the most significantly downregulated gene in lung cancer, aligning with previous studies showing reduced CYP27A1 expression in lung adenocarcinoma [19]. This finding points to substantial perturbation of cholesterol metabolism in the alternative bile acid biosynthesis pathway in lung cancer cells, which we explore in detail in subsequent sections.

In addition, several NUP genes, colored green in Fig 2b, are associated with the DNA transport reaction, which facilitates the movement of DNA between the cytosol and nucleus. These genes demonstrate moderate importance but exhibit high correlations and a prominent distribution among the top-ranked genes. This confirms the involvement of previously reported NUP genes in lung cancer [20,21] while also uncovering novel associations with NUP42,NUP133 and NUP188. These genes, not previously well-established as being linked to lung cancer, were found to be upregulated in cancer samples alongside other known cancer-related nucleoporins shown in Fig 2b.

At the pathway level, our analysis revealed widespread metabolic alterations across multiple subsystems (Fig 2c). The Isolated subsystem, which includes reactions that are not easily categorized into specific metabolic pathways, emerged as the most significant contributor, accounting for 15.1% of the total importance. This was followed by the bile acid biosynthesis pathway, contributing 12.9%, and nucleotide metabolism, which ranked third with 10.8%.

The strength of our integrated approach lies in its ability to connect pathway-level changes with specific gene activities. For instance, the high importance of the bile acid biosynthesis pathway in our analysis was traced back to specific enzymatic reactions and gene expression changes, particularly the downregulation of CYP27A1. Similarly, we identified how alterations in individual genes contributed to broader pathway dysregulation, providing a mechanistic understanding of metabolic reprogramming in lung cancer. These interconnected findings demonstrate the complex nature of metabolic adaptations in lung cancer cells and highlight the importance of analyzing these changes at multiple biological levels simultaneously.

Our multi-level analysis also revealed the significant upregulation of ITPA in lung cancer compared to healthy tissue, aligning with previous study on non-small cell lung cancer [22]. ITPA’s dual involvement in nucleotide and purine metabolism pathways presented an interesting case study for our analytical approach. By simultaneously examining gene expression, reaction fluxes, and pathway relationships, we identified a specific reaction (H2O [c] + XTP [c] ⇒ H+ [c] + PPi [c] + xanthosine-5-phosphate [c]) as particularly crucial in the nucleotide metabolism pathway. While ITPA has traditionally been viewed as a housekeeping gene responsible for nucleotide pool maintenance [23] , our findings suggest a more specialized role in lung cancer metabolism. Beyond its known function in hydrolyzing non-canonical purine nucleotides, the marked upregulation of ITPA appears to serve a critical function in lung cancer cell survival. This adaptation may be particularly important given the elevated demand for purines in rapidly proliferating cancer cells, where ITPA’s enhanced activity helps maintain genome stability by efficiently removing potentially mutagenic non-canonical purines like XTP from the nucleotide pool. In the following sections, we will delve into the biological relevance of the most important findings presented here, providing a detailed explanation.

### Aminoacyl-tRNA biosynthesis and alternative bile acid biosynthesis pathway in lung cancer

Building on the pathway-level insights, we now turn our focus to specific metabolic pathways that exhibit notable differences between lung cancer and healthy lung tissue. In particular, we will examine the aminoacyl-tRNA biosynthesis and bile acid biosynthesis pathways to better understand their distinct roles in lung cancer metabolism. We begin with the less prominent but functionally significant aminoacyl-tRNA biosynthesis pathway, followed by the more extensively altered bile acid biosynthesis pathway.

### Aminoacyl-tRNA biosynthesis in lung cancer

Our multi-level analysis revealed that while all amino acid utilization reactions in the aminoacyl-tRNA biosynthesis pathway are important, four specific amino acids - valine, isoleucine, histidine, and lysine - demonstrated higher importance scores and showed selective upregulation in lung cancer cells. In particular, the reactions corresponding to valine and isoleucine synthesis emerged as highly significant in our RF classifier analysis (Fig 3a and 3d). The synthesis of their corresponding aminoacyl-tRNAs (tRNA-Val and tRNA-Ile) exhibited consistently elevated activity in cancerous tissues compared to healthy samples (Fig 3b and 3e). These scatter plots comparing reaction fluxes between healthy and cancer samples demonstrated a clear upward shift, with a higher density of points appearing above the y=x line, indicating systematically higher flux values in cancer samples. The density distribution plots further supported this pattern, revealing a distinct rightward shift in cancer samples that suggests increased aminoacyl-tRNA synthesis activity. Quantitative analysis of metabolic fluxes (Fig 3c and 3f) showed that median flux values for these aminoacylation reactions were elevated by approximately 15% in cancer cells relative to healthy tissue, representing a 1.15-fold increase.

**Fig 3.**
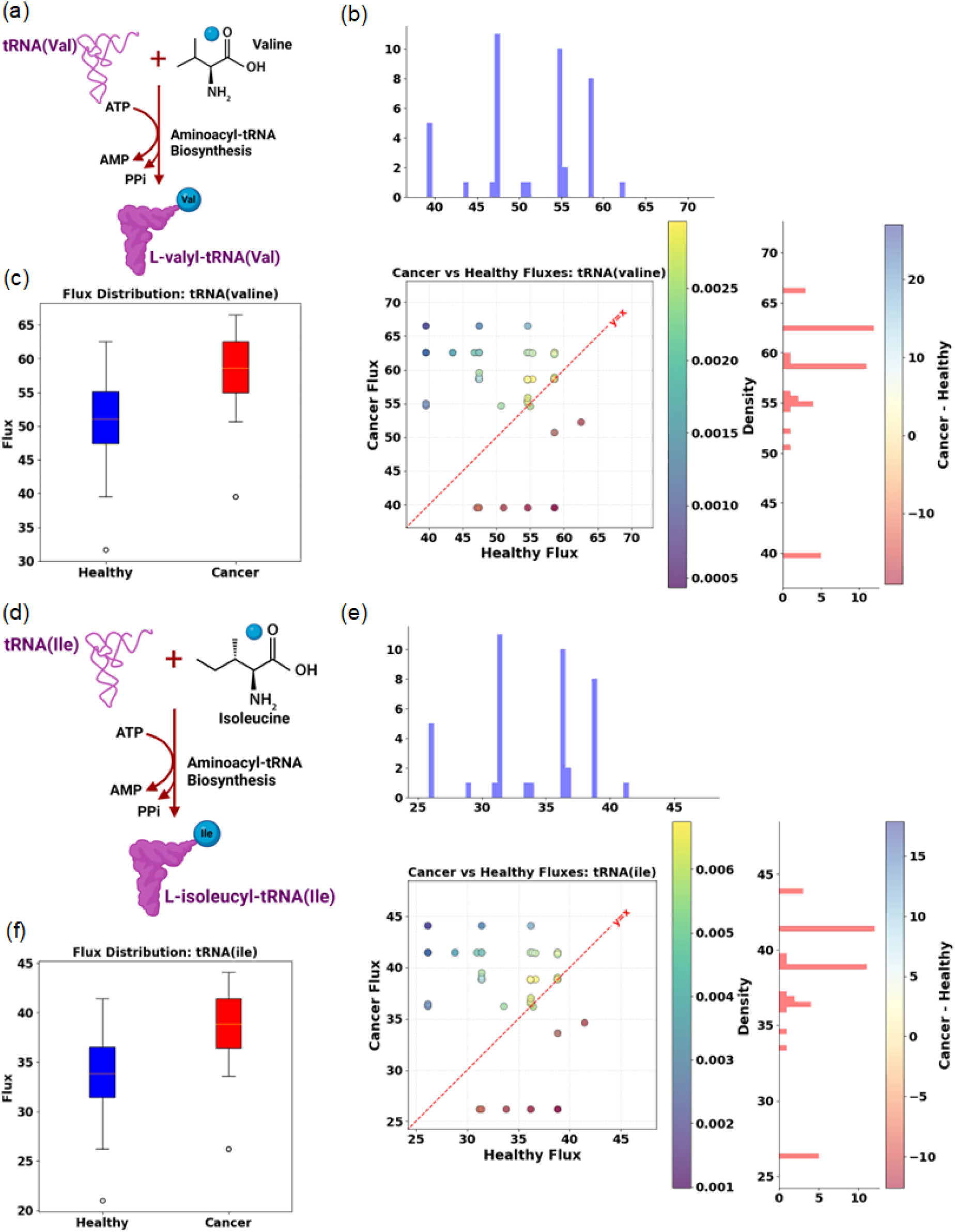
Comparative analysis of aminoacyl-tRNA biosynthesis pathway reactions in lung cancer versus healthy tissue. (a) Schematic representation of the valine aminoacylation reaction (tRNA(Val)), showing the synthesis of L-valyl-tRNA(Val) from valine, ATP, and tRNA. (b) Flux comparison between healthy and cancer samples for the valine aminoacylation reaction, including scatter plot (left) with y=x reference line and density distribution (right and up). (c) Box plot showing the flux distribution comparison between healthy and cancer samples for tRNA(Val) synthesis. (d) Schematic representation of the isoleucine aminoacylation reaction (tRNA(Ile)), showing the synthesis of L-isoleucyl-tRNA(Ile) from isoleucine, ATP, and tRNA. (e) Flux comparison between healthy and cancer samples for the isoleucine aminoacylation reaction, including scatter plot (left) with y=x reference line and density distribution (right and up). (f) Box plot showing the flux distribution comparison between healthy and cancer samples for tRNA(Ile) synthesis. Both reactions demonstrate consistently higher flux values in cancer samples compared to healthy tissue, with approximately 15% elevation in median flux values, indicating increased aminoacyl-tRNA synthesis activity in lung cancer metabolism.

Similar upregulation patterns were observed for histidine and lysine aminoacyl-tRNA synthetase reactions (S4 Fig in supporting information files). While our analysis underscores the importance of all amino acids in the aminoacyl-tRNA pathway, the preferential utilization of these four specific amino acids - valine, isoleucine, histidine, and lysine - suggests a specialized metabolic adaptation in lung cancer cells. This selective upregulation aligns with previous studies that have identified altered branched-chain amino acid metabolism in lung cancer [24,25], but our multi-level approach uniquely reveals the specificity of this adaptation to these four amino acids. The enhanced activity of these particular aminoacyl-tRNA synthesis reactions across the majority of cancer samples points to a precise metabolic reprogramming that may be crucial for supporting the increased energy demands and protein synthesis requirements of rapidly proliferating cancer cells, while also highlighting the overall importance of the entire aminoacyl-tRNA biosynthesis pathway in cancer metabolism.

#### Alternative bile acid biosynthesis in lung cancer

Bile acid biosynthesis proceeds through two distinct pathways: the classical (neutral) pathway and the alternative (acidic) pathway [26]. While the classical pathway accounts for approximately 90% of bile acid synthesis and primarily occurs in the liver [26], our analysis highlights the significance of the alternative pathway in lung cancer. A segment of the bile acid biosynthesis pathway is illustrated in Fig 4a. It demonstrates a specific series of enzymatic reactions that begin in the cytosol and continue into the mitochondria, focusing on the conversion of cholesterol to 3alpha,7alpha,12alpha-trihydroxy-5beta-cholestanate through multiple intermediates. The process initiates with cholesterol in the cytosol, where it undergoes hydroxylation by CH25H (MAR01786) to form 26-hydroxycholesterol. This intermediate is further modified through several enzymatic steps to produce 3alpha,7alpha,12alpha-trihydroxy-5beta-cholestan-26-al. The pathway then proceeds with the action of CYP27A1 (MAR01611) to generate 3alpha,7alpha,12alpha-trihydroxy-5beta-cholestanate.

**Fig 4.**
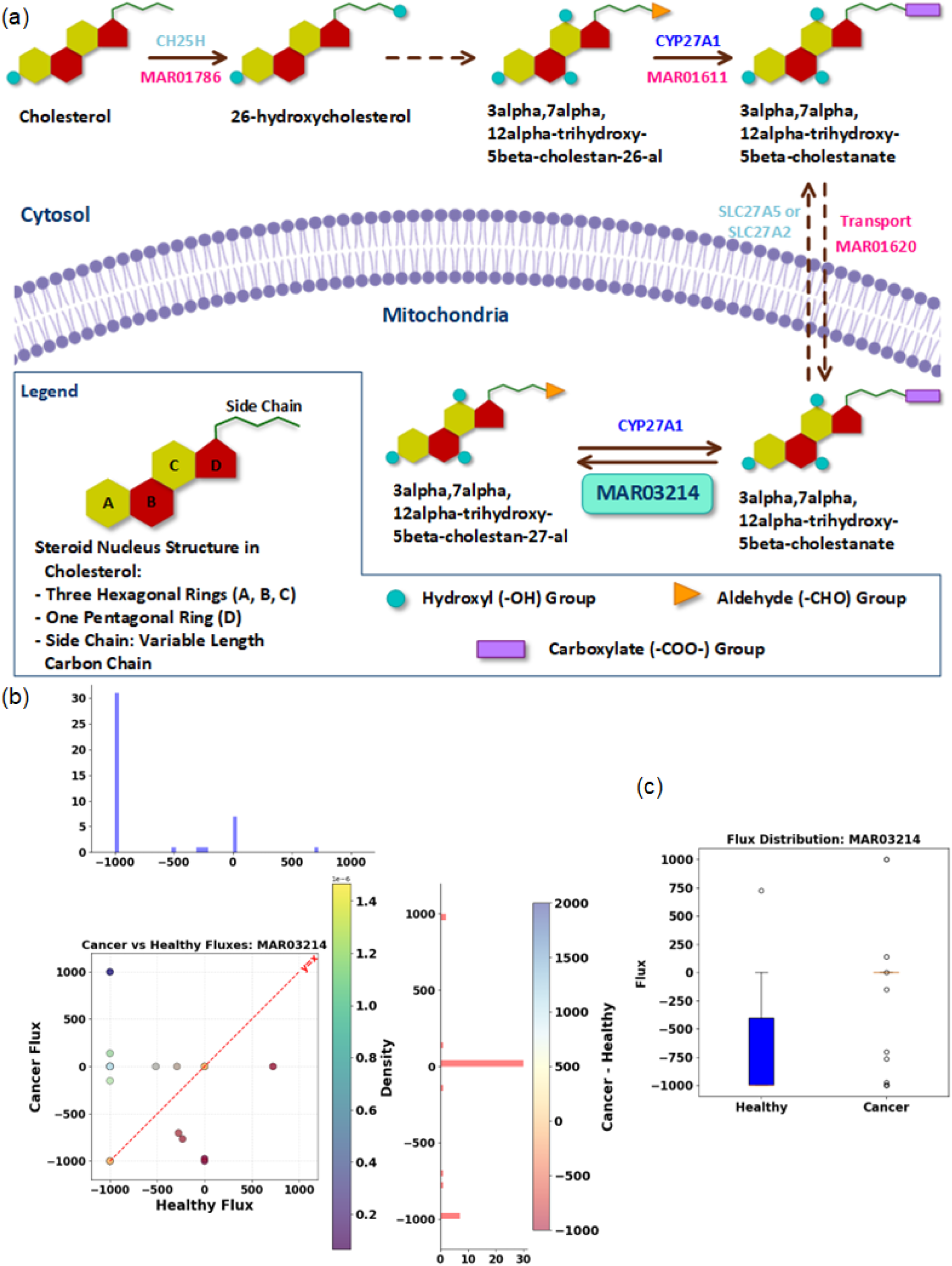
Analysis of CYP27A1-catalyzed reaction (MAR03214) in bile acid biosynthesis pathway. (a) Schematic representation of the bile acid synthesis pathway showing the conversion of cholesterol to bile acids through multiple steps, including the mitochondrial CYP27A1-catalyzed reaction (MAR03214) that interconverts 3alpha,7alpha,12alpha-trihydroxy-5beta-cholestan-27-al and 3alpha,7alpha,12alpha-trihydroxy-5beta-cholestanate. A detailed legend shows the steroid nucleus structure and functional groups. (b) Flux comparison between healthy and cancer samples for reaction MAR03214, including scatter plot with y=x reference line (left) and density distribution (right), showing reduced flux values in cancer samples. (c) Box plot demonstrating the flux distribution comparison between healthy and cancer samples, highlighting significantly lower flux values in cancer tissues.

A critical aspect of this pathway is the transport of metabolites across the mitochondrial membrane, facilitated by specific transporters (SLC27A5 or SLC27A2, MAR01620). Once inside the mitochondria, the pathway culminates in a reversible reaction (MAR03214) catalyzed by CYP27A1, which mediates the interconversion between 3alpha,7alpha,12 alpha-trihydroxy-5beta-cholestan-27-al and 3alpha,7alpha,12 alpha-trihydroxy-5beta-cholestanate. This mitochondrial reaction represents a key step in the alternative (acidic) pathway of bile acid biosynthesis, which, although accounting for only about 10% of total bile acid synthesis, plays a particularly significant role in lung cancer metabolism.

Fig 5a illustrates the other specific segment of the bile acid biosynthesis pathway, focusing on the conversion of cholesterol to 3alpha,7alpha-dihydroxy-5beta-cholestan-26-al through multiple enzymatic steps. The pathway begins with cholesterol, which contains a characteristic steroid nucleus structure comprising three hexagonal rings and one pentagonal ring, along with a variable-length carbon side chain. The initial transformation is catalyzed by CH25H through reaction MAR01784, which introduces a hydroxyl group to form 26-hydroxycholesterol.

**Fig 5.**
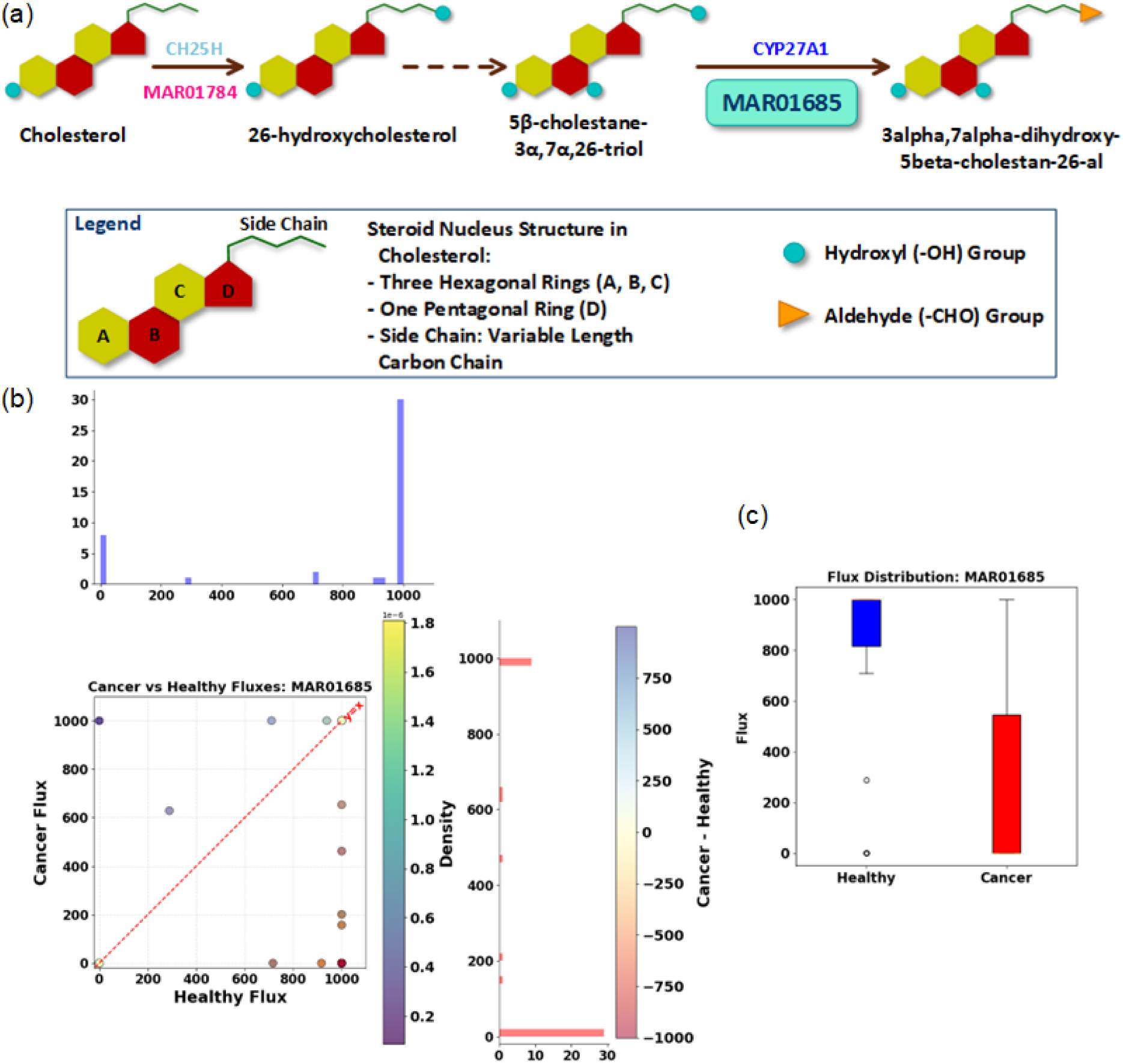
Analysis of the initial CYP27A1-mediated reaction (MAR01685) in the alternative bile acid synthesis pathway. (a) Pathway diagram shows how cholesterol is transformed through multiple enzymatic steps to generate the metabolites involved in reaction MAR01685, which was identified as a key discriminating reaction showing significant downregulation in cancer compared to healthy states. The initial conversion is catalyzed by CH25H (MAR01784), followed by additional enzymatic modifications leading to the metabolites required for the CYP27A1-catalyzed reaction (MAR01685). The legend depicts the fundamental steroid nucleus structure of cholesterol and the functional groups that are attached to this structure during the sequential conversion of cholesterol to pathway intermediates.

Following this initial hydroxylation, the pathway proceeds through an intermediate step to generate 5β-cholestane-3α,7α,26-triol. The final step in this sequence is catalyzed by CYP27A1 through reaction MAR01685, which oxidizes the hydroxyl group at position 26 to form an aldehyde group, resulting in the production of 3alpha,7alpha-dihydroxy-5beta-cholestan-26-al. Throughout these transformations, the steroid nucleus structure remains intact while modifications occur primarily on the side chain and at specific positions of the ring system, as indicated by the presence of hydroxyl (-OH) groups (shown as blue circles) and the final aldehyde (-CHO) group (represented by an orange triangle) in Fig 5a.

Recent studies have demonstrated that alterations in this pulmonary bile acid synthesis pathway, particularly changes in CYP27A1 expression and activity, may be linked to various lung pathologies, including certain types of lung cancer[19]. Our RF classifier identified two key reactions as important discriminating factors in the bile acid biosynthesis pathway, both catalyzed by the CYP27A1 gene (Fig 4a and Fig 5a). The altered activity of CYP27A1 affects the mitochondrial metabolism of cholesterol derivatives, potentially influencing cellular homeostasis and signaling pathways in lung cancer cells.

The flux through reactions catalyzed by CYP27A1 was significantly reduced in lung cancer samples compared to healthy tissues (Fig 4b and Fig 5b). The plots comparing reaction fluxes between healthy and cancer samples demonstrate a clear downward shift, with a higher density of points appearing below the y=x line, indicating systematically lower flux values in cancer samples compared to healthy tissues. This pattern shows two distinct characteristics in the density distribution plots: for positive fluxes, there is a leftward shift in cancer samples indicating reduced activity, while for negative fluxes, cancer samples show a marked shift toward zero flux values compared to the substantial negative fluxes observed in healthy tissues. For reaction MAR03214 (Fig 4b), the flux comparison reveals a consistent pattern of decreased activity in cancer samples, with most data points clustering below the reference line. Similarly, for reaction MAR01685 (Fig 5b), the scatter plot shows a pronounced downward deviation from the y=x line, indicating substantially reduced reaction flux in cancerous tissues. These systematic reductions are quantitatively reflected in the median flux values for these reactions, which were markedly lower in cancer cells compared to healthy tissues (Fig 4c, Fig 5c), suggesting a significant dysregulation of the CYP27A1-mediated bile acid synthesis pathway in lung cancer metabolism.

This decrease in CYP27A1 activity suggests a potential downregulation of bile acid biosynthesis in lung cancer. These reaction flux alterations show how lung cancer cells change their metabolism, particularly in amino acid processing and bile acid production. While the bulk tissue analysis provided valuable insights, we also applied our multi-level approach to the mast cell-specific models to further investigate the role of this kind of immune cell in lung cancer metabolism.

### The role of mast cells in lung cancer

Given the metabolic shifts observed in lung cancer, we next sought to explore how these alterations manifest in specific immune cell populations. Since mast cells play a crucial role in the tumor microenvironment, as evidenced by Fig 1 which demonstrates their high proportion in lung cancer-healthy tissue samples and a notable difference in their distribution between healthy and cancerous tissues, we applied our framework to mast cell-specific metabolic models to further investigate their involvement in lung cancer metabolism. Using reaction fluxes obtained from FBA of the mast cell metabolic models, we implemented a Random Forest classifier to differentiate between healthy and cancerous states. Our analysis revealed significant alterations in mast cell metabolism and function in lung cancer microenvironment. Our RF classifier identified key important reactions in the metabolic network associated with mast cell function in lung cancer. Subsequently, we utilized Gene-Protein-Reaction (GPR) associations, as discussed earlier, related to these reactions to determine the most important genes. This analysis revealed several genes crucial to mast cell function that are significantly altered in the lung cancer state. Fig 6a displays gene importance distribution, where genes are arranged with color intensities ranging from low importance (light yellow) to high importance (dark red), effectively illustrating the varying significance of different genes in mast cell function.

**Fig 6.**
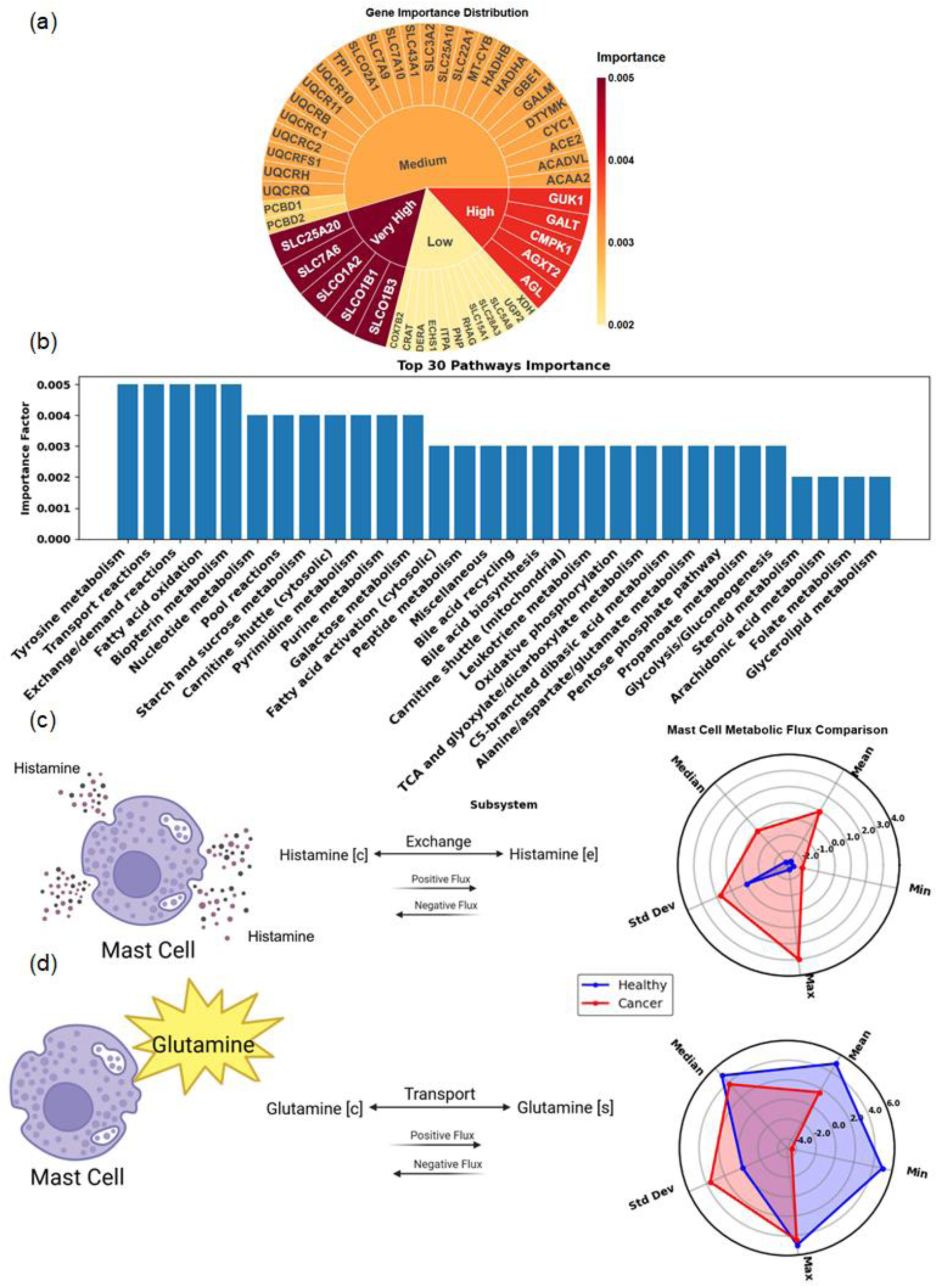
Analysis of mast cell metabolism and metabolic characteristics in lung cancer. (a) Showing the distribution of gene importance in mast cell function, with color intensity indicating significance level (dark red), high (red), medium (orange), and low (light yellow). (b) Ranking the top 30 metabolic pathways by their importance in distinguishing mast cell behavior between normal and lung cancer conditions. (c) Schematic representation of histamine transport dynamics in mast cells, Increased histamine degranulation patterns for mast cells in lung cancer compared to healthy controls, indicating enhanced histamine release into the extracellular space. The plot compares histamine flux pattern between healthy and cancer states (Min, Max, Mean, Standard Deviation of reaction fluxes). (d) Glutamine transport in mast cells with corresponding plot demonstrating metabolic flux differences between normal (blue) and cancer (red) conditions, suggesting increased energy demands during the lung cancer state. The plots in panels (c) and (d) quantitatively represent the differential metabolic patterns observed in normal versus cancerous states, highlighting significant alterations in both histamine and glutamine metabolism.

Based on the gene importance distribution shown in the Fig 6a, the identified genes play distinct roles in mast cell metabolism in lung cancer. The SLCO family members (SLCO1B3, SLCO1B1, SLCO1A2), shown in the very high importance range (dark red section), encode organic anion transporting polypeptides (OATPs) that are crucial for bile acid transport [27]. This transport function is particularly relevant given the role of bile acid biosynthesis in lung cancer metabolism (previous section) and its potential effects on mast cell function in the tumor microenvironment.

SLC7A6, appearing in the very high importance category, encodes the y+LAT2 transporter that is essential for maintaining amino acid homeostasis through its specific transport of cationic amino acids (like L-arginine, L-lysine) and neutral amino acids (such as L-leucine, L-glutamine) [28]. This transport function is critical for protein synthesis and mediator production in mast cells within the lung cancer microenvironment. Similarly, SLC25A20, also in the very high importance category (dark red), functions as a carnitine/acylcarnitine translocase that is fundamental for transporting fatty acids into the mitochondria for beta-oxidation and cellular energy production [29]. These results show that fatty acid oxidation is one of the main energy supplies for mast cell activation and function in the context of lung cancer metabolism.

The identification of these transport-related genes as highly important through the GPR analysis emphasizes how mast cell function in lung cancer is heavily dependent on their ability to maintain proper nutrient exchange and energy metabolism. Their collective high importance suggests that metabolic reprogramming, particularly in terms of amino acid transport and energy generation, is a crucial aspect of mast cell adaptation to the tumor microenvironment.

Fig 6b quantifies the relative importance of different metabolic pathways, with tyrosine metabolism, transport reactions, exchange and demand reactions, and fatty acid oxidation as the most important pathways. The consistent height of several top pathways indicates multiple metabolic processes are simultaneously altered in cancer-associated mast cells. In addition to transport reactions and fatty acid oxidation (with their role in mast cell activation was already explained in the previous section), our analysis revealed tyrosine metabolism alteration, which aligns with previous studies showing its crucial role in mast cell activation. In particular, tyrosine metabolism plays a central role through the KIT receptor signaling pathway in mast cells [30]. Upon stimulation by Stem Cell Factor (SCF), KIT undergoes tyrosine phosphorylation, triggering downstream cascades that activate ERK and AKT kinases [31]. This activation enhances mast cell survival and proliferation [32]. The altered tyrosine metabolism in cancer-associated mast cells leads to enhanced production of inflammatory mediators and bioactive compounds that modify the tumor microenvironment [3]. Additionally, these activated mast cells show increased secretion of proteases like tryptase and chymase, which contribute to extracellular matrix remodeling [33] and can contribute in tumor progression. This metabolic alteration, alongside other pathways shown in Fig 6b, creates an interconnected network that supports mast cell functions and tumor growth and progression in lung cancer, suggesting potential therapeutic targets through tyrosine kinase inhibition.

The most striking functional changes are captured in Fig 6c and 6d, which reveal two key adaptations in the reaction-level analysis. Fig 6c reveals an amplified histamine flux pattern in cancer conditions (red area) compared to normal conditions, with notably higher values across statistical parameters. Specifically, cancer conditions show elevated maximum flux rates and a higher mean value, while maintaining a broader range between minimum and maximum values. The increased standard deviation in cancer conditions suggests more variable histamine flux dynamics, while the median values indicate that this enhanced histamine transport is sustained rather than driven by occasional spikes. This enhanced histamine transport suggests mast cells become more actively degranulating in the lung cancer environment.

The metabolic shift is further highlighted in Fig 6d, where the radar plot compares glutamine transport parameters between healthy and cancer conditions. The healthy cases exhibit relatively balanced values, with the minimum and maximum being roughly similar and a standard deviation close to zero. In contrast, the cancer cases display a negative minimum, a positive maximum, and a significantly larger standard deviation. The shift of glutamine transport reaction fluxes towards negative in cancer conditions likely reflects the enhanced glutamine uptake required to meet the heightened energy demands of mast cells within the tumor microenvironment.

These findings collectively demonstrate the complex metabolic adaptations of mast cells in lung cancer and highlight their active participation in shaping the tumor microenvironment through both metabolic reprogramming and enhanced degranulation responses.

### Temperature-dependent metabolic analysis

Research has demonstrated that temperature plays a significant role in cancer therapy, particularly in enhancing immune responses against tumors [34]. This insight into the thermal sensitivity of cancer cells and immune functions has motivated further investigations into temperature effects on the growth rates of both cancer cells and immune cells, such as mast cells, in various microenvironments including the lung. Given the extensive reaction network in Human1 and its intricate pathway interactions, we developed the Metabolic Temperature Sensitivity Analysis (MTSA) method to specifically investigate temperature effects on both cancer cell growth and mast cell behavior in the lung cancer microenvironment. This approach is particularly valuable as it provides insights into how temperature variations affect the complex interplay between lung cancer metabolism and mast cell function. Our analysis using MTSA revealed distinct biomass distribution patterns between healthy and cancerous conditions in both lung tissue and mast cells within the lung cancer microenvironment (Fig 7 and Fig 8). We investigated the clinically significant temperature spectrum of 36-40°C, which spans from normal body temperature (37°C) to fever states frequently observed in cancer patients. The plots demonstrate a clear bimodal distribution in biomass production across this temperature range, with notable differences between healthy and diseased states.

**Fig 7:**
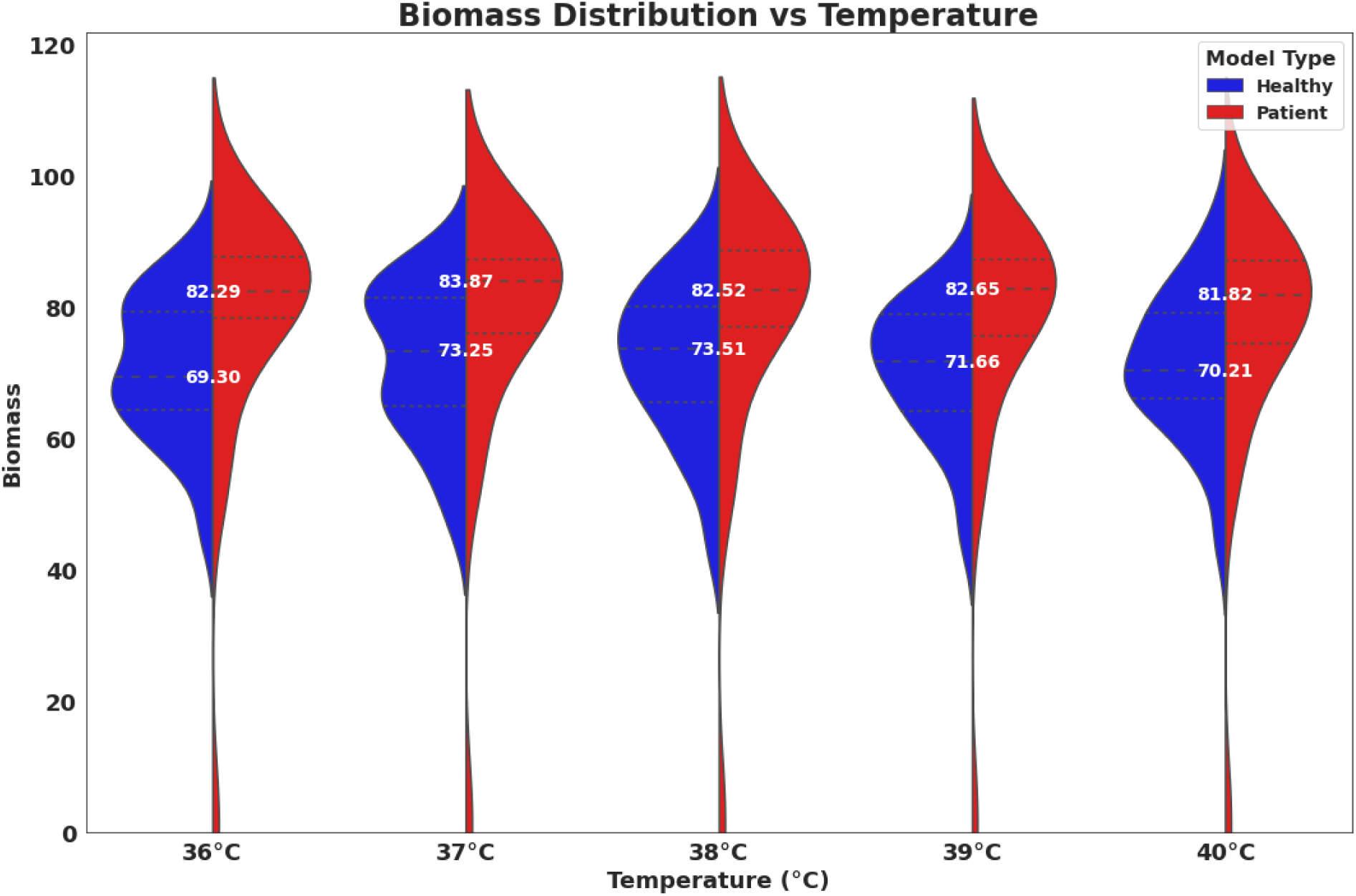
Tissue-level biomass distribution analysis comparing healthy and cancerous lung tissue across temperature range 36-40°C. The plots demonstrate that cancerous tissue (red) maintains higher mean biomass values compared to healthy tissue (blue), with peak production occurring at different optimal temperatures - 37°C for cancerous tissue (83.87) and 38°C for healthy tissue (82.52), suggesting distinct temperature-dependent metabolic preferences between the two conditions. The distributions demonstrate a gradual decline in biomass production as temperature increases, with a notable decrease at 40°C for both tissue types.

**Figure 8:**
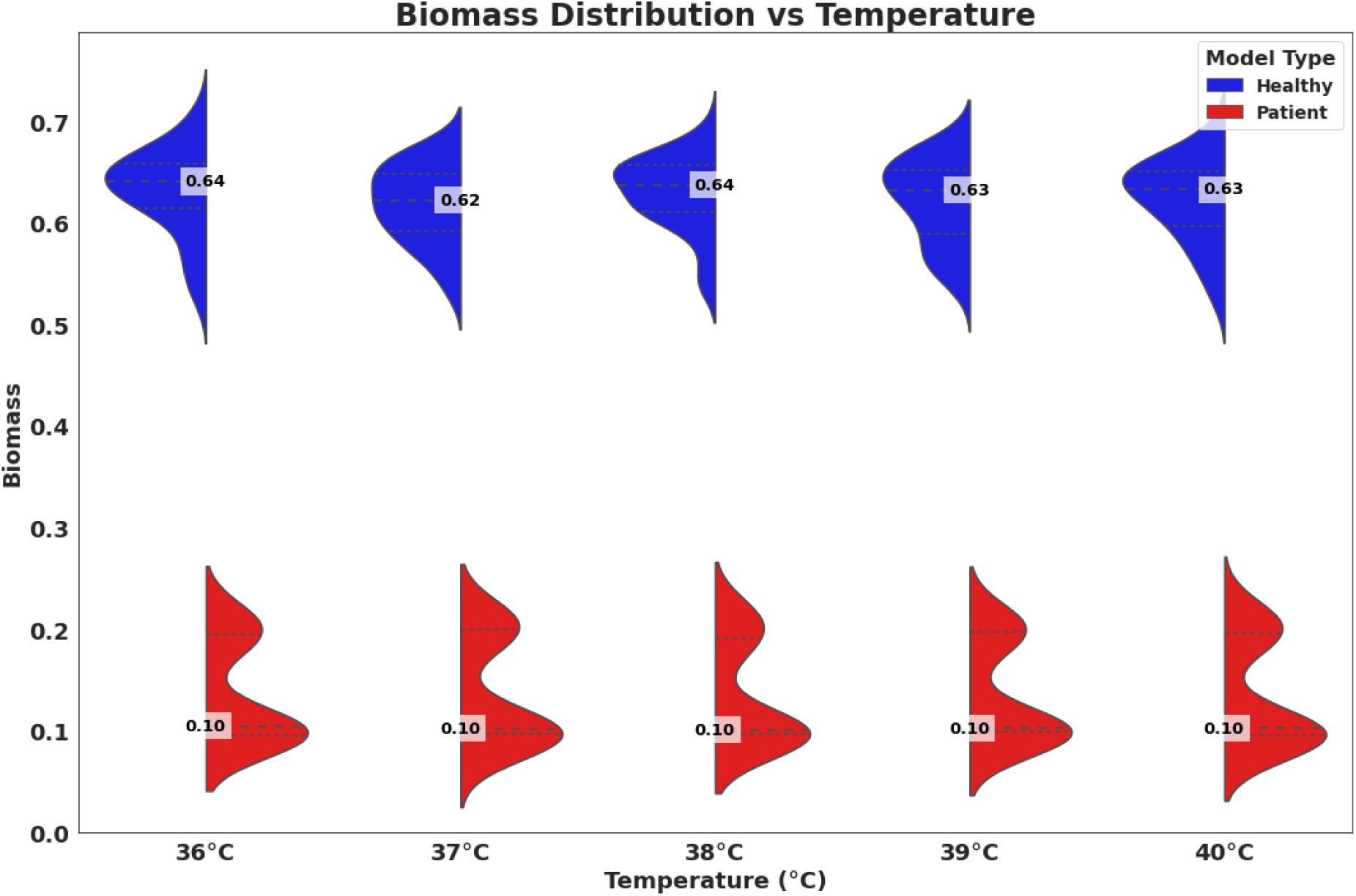
Temperature-dependent biomass distribution in healthy and cancerous mast cells analyzed using the MTSA method. The plots show the contrasting biomass patterns between healthy (blue) and cancerous (red) mast cells across temperatures from 36°C to 40°C. Healthy mast cells maintained high biomass values (0.62-0.64) with slight variations across temperatures, showing particular resilience at fever-range temperatures (38-40°C). In contrast, cancerous mast cells maintained a consistently low biomass (0.10) across all temperature points, suggesting that the lung cancer microenvironment significantly suppresses mast cell growth and metabolic activity. This dramatic reduction in biomass indicates that the tumor microenvironment creates conditions that substantially impair mast cell proliferation.

In lung tissue samples, cancerous tissue consistently demonstrated higher mean biomass values compared to healthy tissue across all temperature points (Fig 7). Healthy tissues (blue) exhibited optimal biomass production at 38°C, while patient tissues showed peak biomass at 37°C. As temperatures increased beyond these optimal points, both tissue types exhibited a gradual decrease in biomass production. We found that both tissue types exhibited their lowest biomass production at 40°C, with cancerous tissues showing a steeper reduction than healthy tissues.

The analysis of mast cell populations revealed a striking contrast between healthy and cancerous cells across different temperatures (Fig 8). Healthy mast cells exhibited significantly higher mean biomass values compared to their cancerous counterparts at all temperature points. In healthy mast cells, the mean biomass started at approximately 0.64 at 36°C, dipped slightly to 0.62 at 37°C, and then recovered to stabilize around 0.63–0.64 in the fever range (38–39°C). At 40°C, the biomass remained steady at 0.63, demonstrating remarkable thermal resilience. In contrast, cancerous mast cells consistently displayed a mean biomass of approximately 0.10 across all temperatures, with minimal variation in response to temperature changes. This stark difference suggests that the tumor microenvironment substantially inhibits mast cell proliferation and metabolic activity, likely through alterations in signaling pathways or resource availability. The robust thermal response observed in healthy mast cells, particularly in the fever range, may reflect an evolutionary adaptation linked to their immune function. However, this adaptive capability appears to be lost in the cancerous state, as cancerous mast cells fail to exhibit significant biomass changes across temperatures. These findings underscore the potential influence of thermal regulation on mast cell function and its broader implications for cancer progression.

## Discussion

Our study presents key contributions to understanding lung cancer metabolism through the development of genome-scale metabolic models for both lung tissue and mast cells in the context of cancer. These comprehensive models, built upon the extensive Human1 framework, provide unique insights into the metabolic reprogramming that occurs in lung cancer and its immune microenvironment. Our multi-level approach, which integrates gene expression, reaction fluxes, and pathway analysis, offers a powerful platform for investigating the complex metabolic interactions between cancer cells and immune components.

Our cell-type deconvolution analysis revealed a significant reduction in resting mast cells within cancerous tissues, providing the foundation for our subsequent metabolic investigations. The analysis demonstrated substantial alterations in immune cell populations, particularly highlighting the critical role of mast cells in the tumor microenvironment. This finding motivated our development of specialized metabolic models to understand the functional implications of these cellular changes.

The machine learning analysis of our metabolic models, combined with our multi-level analytical approach, revealed critical alterations in several key pathways. Most significantly, we discovered a specialized metabolic adaptation in lung cancer cells involving the selective upregulation of four specific amino acids (valine, isoleucine, histidine, and lysine) in the aminoacyl-tRNA pathway. The synthesis reactions for these amino acids showed approximately 15% higher flux values in cancerous tissues compared to healthy tissue. This preferential utilization suggests a precise metabolic strategy employed by cancer cells to meet their elevated energy requirements. By simultaneously analyzing gene, reaction and pathway importances, we were able to identify not only which amino acids were upregulated but also understand the broader context of how this selective usage supports cancer cell proliferation and survival. In addition, our multi-level analysis not only revealed previously known dysregulated genes, reactions and pathways in lung cancer, but also discovered new important dysregulations like NUP42, NUP133, and NUP188 genes functions, which are not well studied in lung cancer.

The subsequent analysis of mast cell metabolism in the tumor microenvironment revealed striking adaptations, including enhanced histamine transport and increased glutamine consumption. These changes suggest a shift toward a more active state in cancer-associated mast cells. The identification of key transport-related genes, including the SLCO family members and SLC transporters, highlights the critical role of metabolite transport in mast cell function within the tumor environment. These findings provide new perspectives on how mast cells adapt their metabolism to support or potentially inhibit tumor progression.

A major methodological advancement of our study is the development of the Metabolic Thermodynamic Sensitivity Analysis (MTSA) method, which addresses a critical gap in cancer metabolism research. By integrating thermodynamic principles with genome-scale metabolic modeling, MTSA enables systematic investigation of temperature-dependent metabolic changes. This approach is particularly significant, as it allows for the quantitative prediction of how thermal fluctuations—arising from fever, inflammation, or therapeutic interventions—may influence both cancer progression and immune cell function. The ability of MTSA to analyze multiple metabolic pathways while accounting for temperature effects provides a powerful framework for studying the intricate relationship between thermal conditions and cellular metabolism in cancer. This innovative approach has broad applicability beyond the scope of our current study. The MTSA method can be readily adapted to investigate metabolic responses to temperature changes in various fields of biology, including different organisms, other cancer types, and diverse immune cell populations. Its versatility makes it a valuable tool for exploring temperature-dependent metabolic phenomena in fields such as immunology, microbiology, plant biology, and environmental science. Applying MTSA to our models revealed distinct temperature-dependent metabolic patterns that differentiate cancerous from healthy states. Notably, while healthy lung tissue exhibited peak biomass production at 38°C, cancerous tissue maintained higher biomass production at 37°C, suggesting fundamental differences in temperature-dependent metabolic regulation. More strikingly, cancer-associated mast cells showed a dramatic reduction in biomass production across all temperatures (36–40°C), indicating a profound metabolic impairment in these immune cells within the tumor microenvironment. These findings highlight the potential impact of temperature fluctuations on both tumor progression and immune function.

These discoveries open several promising avenues for future research and therapeutic interventions. The selective upregulation of specific aminoacyl-tRNA synthesis pathways suggests potential therapeutic targets, particularly in combination with existing treatments that disrupt protein synthesis in cancer cells. Additionally, the temperature-dependent metabolic patterns identified through MTSA highlight the potential for temperature-based therapeutic strategies, which could influence the balance between cancer cell growth and immune cell function.

Our genome-scale metabolic models and the MTSA method provide valuable tools for advancing cancer metabolism research. While our computational approach offers comprehensive metabolic insights, it relies on steady-state assumptions that may not fully capture the dynamic nature of cancer metabolism. Additionally, the use of bulk tissue data, despite our deconvolution methods, may not completely account for cellular heterogeneity within the tumor microenvironment. Nevertheless, our multi-level approach successfully identified meaningful metabolic signatures that warrant further investigation. Future experimental validation, particularly focusing on the selective amino acid utilization patterns and temperature-dependent metabolic adaptations of both cancer cells and mast cells, could help overcome these limitations and pave the way for novel therapeutic strategies. Furthermore, while the metabolic biomarkers identified in this study show promise for enhancing early detection and monitoring of lung cancer progression, their clinical utility needs to be validated in larger patient cohorts.

Through the integration of genome-scale metabolic modeling and advanced computational techniques, our study provides novel insights into lung cancer metabolism and the role of mast cells in the tumor microenvironment. Our multi-level approach has revealed previously unknown metabolic adaptations, particularly in amino acid utilization, while the introduction of MTSA has opened new possibilities for understanding temperature-dependent metabolic regulation in cancer.

## Methods

### Data acquisition and preprocessing

Gene expression data from 43 pairs of lung tissue samples were obtained from the GEO dataset (GSE18842). After acquiring the gene expressions, we mapped the genes to those in the Human1 [15] model, resulting in a final set of 3,502 genes with metabolic functions and their corresponding expression values.

### Cell type deconvolution analysis

To investigate mast cell-specific gene expression within our heterogeneous tissue samples, we employed CIBERSORTx [14], a powerful machine learning tool developed for assessing cellular abundance and cell type-specific gene expression patterns from bulk tissue transcriptome profiles. CIBERSORTx allows for the deconvolution of complex gene expression mixtures, enabling the estimation of cell type proportions and gene expression profiles without the need for physical cell isolation [14].

We utilized CIBERSORTx to impute mast cell-related marker gene expressions from the bulk RNA-seq data for all healthy and cancerous samples in our dataset. This deconvolution method enabled us to estimate the gene expression profiles of mast cells within the heterogeneous tissue samples.

### Genome-scale metabolic modeling

We developed genome-scale metabolic models using the Human1 model and the iMAT [16] method. First, Genes were mapped to their corresponding reactions based on gene-protein-reaction (GPR) associations. Reaction expression levels were then calculated using these gene-to-reaction mappings and the established GPR rules. Reaction expression levels were categorized into three groups: lowly expressed, moderately expressed, and highly expressed. Based on the characteristics of the dataset, two thresholds were established for this categorization: a lower threshold defined as the mean expression value minus half of the standard deviation (Mean - 0.5 * standard deviation (STD)), and an upper threshold defined as the mean expression value plus half of the standard deviation (Mean + 0.5 * standard deviation (STD)). Reactions with expression levels below the lower threshold were classified as lowly expressed, those above the upper threshold were considered highly expressed, and the reactions with expression levels falling between these two thresholds were categorized as moderately expressed. We chose 0.5 * SD based on the characteristics of our dataset, as it provided a suitable distribution of reaction expression across the three expression categories.

Afterwards, iMAT was used to focus on highly expressed reaction, and to exclude lowly expressed reactions, which often have higher variability [35], from our analysis. This filtering strategy enhances computational efficiency by reducing the number of models’ reactions. In addition, the highly expressed reactions are likely to have a more significant impact on cellular functions and metabolic processes. For each sample, we generated two types of models: lung bulk tissue-specific models and lung-associated mast cells models for all samples. For the lung-associated mast cells models, we used a biomass equation similar to that of macrophages [36] in lung tissue. This approach was necessitated by the lack of a specific biomass equation for mast cells in the literature. Additionally, this choice was supported by several shared characteristics between mast cells and macrophages. Both cell types share a myeloid lineage, indicating a common origin [37], and exhibit functional similarities in lung tissue, such as roles in immune response and inflammation [38]. Furthermore, both mast cells and macrophages contain granular content [39,40], which is crucial for their immune defense roles. These similarities justify the use of the macrophage biomass equation as a proxy for mast cells, enabling us to proceed with our analysis despite the lack of mast cell-specific data. Finally, Flux Balance Analysis (FBA) [41] was performed to obtain the reaction flux profiles for each sample. FBA is a widely used computational method in systems biology that calculates the flow of metabolites through a metabolic network, predicting the growth rate of an organism or the rate of production of an important metabolite. It uses linear programming to determine a set of metabolic fluxes that optimize a given objective function, such as biomass production, while satisfying mass balance constraint. In our study, FBA allowed us to predict the metabolic capabilities of our lung tissue and lung - associated mast cell models under steady-state conditions, providing insights into the metabolic differences between normal and cancerous states.

### Machine learning analysis

#### Random forest classifiers for healthy and cancerous case categorizations

We employed a Random Forest classifier to distinguish between healthy and cancerous samples using reaction flux data from FBA as the input. The dataset was split into training (80%) and test (20%) set, with stratification to ensure balanced class representation. The Random Forest model was implemented using scikit-learn with 1000 trees, Gini impurity criterion, and a maximum depth of 32. The model’s performance was assessed using 8-fold cross-validation and out-of-bag (OOB) score estimation. Feature importance scores were utilized to identify the most discriminating reactions between healthy and cancerous samples.

### Temperature effect analysis: Metabolic Thermodynamic Sensitivity Analysis (MTSA)

We developed a novel method for thermodynamics-based calculation of the effect of temperature changes on cancer growth rate and mast cell growth rate. This method, which we call Metabolic Thermodynamic Sensitivity Analysis (MTSA), integrates various computational approaches to provide a comprehensive understanding of temperature-dependent metabolic changes. Due to the complexity of the models and limitations in kinetic data availability, we made several key assumptions. We assumed that all enzymatic reaction rates follow the Michaelis-Menten equation [42]:

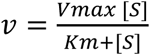

where *v* is the reaction rate, *V_max_* is the maximum reaction rate, [*S*] is the substrate concentration, and *K_m_* is the Michaelis constant. Additionally, we assumed that each reaction operates at maximum driving force, as all reactions are in pseudo-steady state and occur rapidly. Furthermore, the reaction rate is calculated based on the rate-limiting substrate. Among all the substrates of a reaction, the one having the minimum value of *k_cat_*/*K_m_* is considered as the rate-limiting substrate. We used the Max/Min Driving Force (MDF) method [43] to calculate metabolite concentrations that maximize the Gibbs free energy of each reaction. To find the rate of each reaction within the feasible solution space, we employed Flux Variability Analysis (FVA) [44] to determine the maximum flux for each reaction which served as a reference point for generating a range of potential *V_max_* values. The *k_cat_* values for each enzyme were estimated using the DL-Kcat method [45]. Through non-linear regression, we determined the Michaelis constant and reaction rate within the feasible space for the rate-limiting substrate of each reaction. This approach allowed for the robust estimation of kinetic parameters while accounting for the inherent nonlinearity of enzyme-catalyzed reactions (See S1 text in supporting information for more details). This comprehensive methodology, MTSA, allowed us to investigate the effects of temperature on cancer and mast cell growth rates.

## Acknowledgments

We gratefully acknowledge the Holland Computing Center (HCC) at the University of Nebraska-Lincoln for providing high-performance computing resources and technical support that contributed to the results presented in this research.

## Data availability

All code used to reproduce this publication’s data, and the metabolic models are available at our GitHub repository: https://github.com/ssbio/MTSA.git

## Supporting information captions

**S1 Fig. The distribution of reactions and metabolites across healthy lung tissue models.**

**S2 Fig. The distribution of reactions and metabolites across cancerous lung tissue models.**

**S1 Table. Performance comparison of RF classifiers for lung bulk tissue and mast cell metabolic models.**

**S3 Fig. Analysis of metabolic flux distributions in MAR08643 reaction**

**S4 Fig. Analysis of metabolic flux distributions in aminoacyl-tRNA synthesis pathway between cancer and healthy lung tissues.**

**S1 text. MTSA method description**

## Notes

### Competing Interest Statement

The authors have declared no competing interest.

## References

1. Li C, Lei S, Ding L, Xu Y, Wu X, Wang H, Zhang Z, Gao T, Zhang Y, Li L. Global burden and trends of lung cancer incidence and mortality. Chin Med J (Engl). 2023;136(13):1583–90.

2. Herath S, Sadeghi Rad H, Radfar P, Ladwa R, Warkiani M, O’Byrne K, Kulasinghe A. The role of circulating biomarkers in lung cancer. Front Oncol. 2022;11:801269.

3. Komi DEA, Mortaz E, Amani S, Tiotiu A, Folkerts G, Adcock IM. The role of mast cells in IgE-independent lung diseases. Clin Rev Allergy Immunol. 2020;58:377–87.

4. Imada A, Shijubo N, Kojima H, Abe S. Mast cells correlate with angiogenesis and poor outcome in stage I lung adenocarcinoma. Eur Respir J. 2000;15(6):1087–93.

5. Oldford SA, Marshall JS. Mast cells as targets for immunotherapy of solid tumors. Mol Immunol. 2015;63(1):113–24.

6. Maltby S, Khazaie K, McNagny KM. Mast cells in tumor growth: angiogenesis, tissue remodelling and immune-modulation. Biochim Biophys Acta BBA-Rev Cancer. 2009;1796(1):19–26.

7. Batta I, Patial R, Sobti RC, Agrawal DK. Computational Biology in the Discovery of Biomarkers in the Diagnosis, Treatment and Management of Cardiovascular Diseases. Cardiol Cardiovasc Med. 2024;8(5):405.

8. Passi A, Tibocha-Bonilla JD, Kumar M, Tec-Campos D, Zengler K, Zuniga C. Genome-scale metabolic modeling enables in-depth understanding of big data. Metabolites. 2021;12(1):14.

9. Islam MM, Goertzen A, Singh PK, Saha R. Exploring the metabolic landscape of pancreatic ductal adenocarcinoma cells using genome-scale metabolic modeling. Iscience. 2022;25(6).

10. Baloni P, Dinalankara W, Earls JC, Knijnenburg TA, Geman D, Marchionni L, Price ND. Identifying personalized metabolic signatures in breast cancer. Metabolites. 2020;11(1):20.

11. Zhang C, Aldrees M, Arif M, Li X, Mardinoglu A, Aziz MA. Elucidating the reprograming of colorectal cancer metabolism using genome-scale metabolic modeling. Front Oncol. 2019;9:681.

12. Larsson I, Uhlén M, Zhang C, Mardinoglu A. Genome-scale metabolic modeling of glioblastoma reveals promising targets for drug development. Front Genet. 2020;11:381.

13. Zhang B, Shi H, Wang H. Machine learning and AI in cancer prognosis, prediction, and treatment selection: a critical approach. J Multidiscip Healthc. 2023;1779–91.

14. Newman AM, Steen CB, Liu CL, Gentles AJ, Chaudhuri AA, Scherer F, Khodadoust MS, Esfahani MS, Luca BA, Steiner D. Determining cell type abundance and expression from bulk tissues with digital cytometry. Nat Biotechnol. 2019;37(7):773–82.

15. Robinson JL, Kocabaş P, Wang H, Cholley PE, Cook D, Nilsson A, Anton M, Ferreira R, Domenzain I, Billa V. An atlas of human metabolism. Sci Signal. 2020;13(624):eaaz1482.

16. Zur H, Ruppin E, Shlomi T. iMAT: an integrative metabolic analysis tool. Bioinformatics. 2010;26(24):3140–2.

17. Son S, Stevens MM, Chao HX, Thoreen C, Hosios AM, Schweitzer LD, Weng Y, Wood K, Sabatini D, Vander Heiden MG. Cooperative nutrient accumulation sustains growth of mammalian cells. Sci Rep. 2015;5(1):17401.

18. Yilmaz B, Karabay AZ. Food additive sodium benzoate (NaB) activates NFκB and induces apoptosis in HCT116 cells. Molecules. 2018;23(4):723.

19. Yin Y, He M, Huang Y, Xie X. Transcriptomic analysis identifies CYP27A1 as a diagnostic marker for the prognosis and immunity in lung adenocarcinoma. BMC Immunol. 2023;24(1):37.

20. Nong J, Zhou X, Liu J, Luo J, Huang J, Xie H, Yang K, Wang J, Ye X, Peng T. Nucleoporin 107 is a prognostic biomarker in hepatocellular carcinoma associated with immune infiltration. Cancer Med. 2023;12(9):10990–1009.

21. Fujitomo T, Daigo Y, Matsuda K, Ueda K, Nakamura Y. Critical function for nuclear envelope protein TMEM209 in human pulmonary carcinogenesis. Cancer Res. 2012;72(16):4110–8.

22. Mamoor S. The phosphatase ITPA is differentially expressed in non-small cell lung cancer and its expression associates with human survival. 2024.

23. Simone PD, Struble LR, Kellezi A, Brown CA, Grabow CE, Khutsishvili I, Marky LA, Pavlov YI, Borgstahl GE. The human ITPA polymorphic variant P32T is destabilized by the unpacking of the hydrophobic core. J Struct Biol. 2013;182(3):197–208.

24. Bian M, Huang S, Yu D, Zhou Z. tRNA metabolism and lung cancer: beyond translation. Front Mol Biosci. 2021;8:659388.

25. Sivanand S, Vander Heiden MG. Emerging roles for branched-chain amino acid metabolism in cancer. Cancer Cell. 2020;37(2):147–56.

26. Smith LP, Nierstenhoefer M, Yoo SW, Penzias AS, Tobiasch E, Usheva A. The bile acid synthesis pathway is present and functional in the human ovary. PLoS One. 2009;4(10):e7333.

27. Schulte RR, Ho RH. Organic anion transporting polypeptides: emerging roles in cancer pharmacology. Mol Pharmacol. 2019;95(5):490–506.

28. Bhutia YD, Ganapathy V. Glutamine transporters in mammalian cells and their functions in physiology and cancer. Biochim Biophys Acta BBA-Mol Cell Res. 2016;1863(10):2531–9.

29. Tonazzi A, Giangregorio N, Console L, Palmieri F, Indiveri C. The mitochondrial carnitine acyl-carnitine carrier (SLC25A20): molecular mechanisms of transport, role in redox sensing and interaction with drugs. Biomolecules. 2021;11(4):521.

30. Longo V, Catino A, Montrone Mi, Galetta D, Ribatti D. Controversial role of mast cells in NSCLC tumor progression and angiogenesis. Thorac Cancer. 2022;13(21):2929–34.

31. Möller C, Alfredsson J, Engström M, Wootz H, Xiang Z, Lennartsson J, Jönsson JI, Nilsson G. Stem cell factor promotes mast cell survival via inactivation of FOXO3a-mediated transcriptional induction and MEK-regulated phosphorylation of the proapoptotic protein Bim. Blood. 2005;106(4):1330–6.

32. Shemesh R, Laufer-Geva S, Gorzalczany Y, Anoze A, Sagi-Eisenberg R, Peled N, Roisman LC. The interaction of mast cells with membranes from lung cancer cells induces the release of extracellular vesicles with a unique miRNA signature. Sci Rep. 2023;13(1):21544.

33. Zhao XO, Sommerhoff CP, Paivandy A, Pejler G. Mast cell chymase regulates extracellular matrix remodeling-related events in primary human small airway epithelial cells. J Allergy Clin Immunol. 2022;150(6):1534–44.

34. Repasky EA, Evans SS, Dewhirst MW. Temperature matters! And why it should matter to tumor immunologists. Cancer Immunol Res. 2013;1(4):210–6.

35. Eling N, Richard AC, Richardson S, Marioni JC, Vallejos CA. Correcting the mean-variance dependency for differential variability testing using single-cell RNA sequencing data. Cell Syst. 2018;7(3):284–94.

36. Bordbar A, Lewis NE, Schellenberger J, Palsson BØ, Jamshidi N. Insight into human alveolar macrophage and M. tuberculosis interactions via metabolic reconstructions. Mol Syst Biol. 2010;6(1):422.

37. Galli SJ, Borregaard N, Wynn TA. Phenotypic and functional plasticity of cells of innate immunity: macrophages, mast cells and neutrophils. Nat Immunol. 2011;12(11):1035–44.

38. Chen L, Deng H, Cui H, Fang J, Zuo Z, Deng J, Li Y, Wang X, Zhao L. Inflammatory responses and inflammation-associated diseases in organs. Oncotarget. 2017;9(6):7204.

39. Tan BH, Meinken C, Bastian M, Bruns H, Legaspi A, Ochoa MT, Krutzik SR, Bloom BR, Ganz T, Modlin RL. Macrophages acquire neutrophil granules for antimicrobial activity against intracellular pathogens. J Immunol. 2006;177(3):1864–71.

40. Amin K. The role of mast cells in allergic inflammation. Respir Med. 2012;106(1):9–14.

41. Orth JD, Thiele I, Palsson BØ. What is flux balance analysis? Nat Biotechnol. 2010;28(3):245–8.

42. Michaelis L, Menten ML. Die kinetik der invertinwirkung. Biochem Z. 1913;49(333– 369):352.

43. Noor E, Bar-Even A, Flamholz A, Lubling Y, Davidi D, Milo R. An integrated open framework for thermodynamics of reactions that combines accuracy and coverage. Bioinformatics. 2012;28(15):2037–44.

44. Mahadevan R, Schilling CH. The effects of alternate optimal solutions in constraint-based genome-scale metabolic models. Metab Eng. 2003;5(4):264–76.

45. Li F, Yuan L, Lu H, Li G, Chen Y, Engqvist MK, Kerkhoven EJ, Nielsen J. Deep learning-based k cat prediction enables improved enzyme-constrained model reconstruction. Nat Catal. 2022;5(8):662–72.

